# A foundational *in vivo* platform for predicting human health outcomes

**DOI:** 10.64898/2026.08.03.742611

**Authors:** Thomas Roseberry, Tim Krausz, Gina Williams, David Tingley

## Abstract

Rodents remain the workhorse of preclinical drug development, yet often fail to predict human clinical outcomes. Existing alternatives are similarly constrained. Cells in culture cannot recapitulate whole-organism physiology, and larger mammals cannot be studied at comparable throughput. Here we present a scalable, information-dense platform that can predict a drug’s long-term human clinical outcomes from 24 hours of rodent behavior. A novel home-cage system continuously records behavior, generating thousands of features per hour. Models are trained on human clinical trial data to map these features onto outcomes including gastrointestinal adverse events, cardiac toxicity, neuropsychiatric side effects, and long-term weight loss. In addition to being an order of magnitude faster, the platform provides more accurate clinical predictions than standard long-term preclinical experiments. The approach readily extends to other outcomes, enabling rodents to serve as quantitative models for human clinical prediction.

## Introduction

Roughly one in ten drug candidates that enters a clinical trial reaches approval (Hay et al. 2014; Wong et al. 2019). Among programs that fail for an identified reason, the largest share comes from failures in efficacy or toxicity in humans (Harrison 2016). Contemporary estimates place the capitalized cost of bringing a single new drug to market well over one billion dollars, a figure dominated by the price of failure (DiMasi et al. 2016; Paul et al. 2010). Compounding this cost, a drug needs to be a blockbuster to return the total investment and cover the costs of failures and non-blockbusters (Fernandez et al. 2012). Preclinical animal experiments are meant to de-risk this process, yet they have historically been poor predictors of a drug’s performance (Bracken 2009).

Driving this failure is a problem with how animal experiments are conceived and conducted. A conventional preclinical study reduces a drug’s whole-body effect to one or a few pre-selected readouts, each chosen because the experimenter expects it to stand in for the human clinical state. However, mice are not humans and the approach lacks clinical predictive validity (Garner 2014; Dirnagl 2021). For example, mice cannot report pain. Analgesic efficacy is often inferred from evoked withdrawal reflexes that have face validity but poor predictive validity, and compounds that reverse this proxy in rodents have repeatedly failed to relieve pain in patients (Mogil 2009; Taneja et al. 2012).

Practical limits on *in vivo* data production are set by time and money rather than biological questions. While roughly 110 million mice and rats are used in research each year (Carbone et al. 2021), the core workflow, in which an experimenter manually handles, doses, and observes individual animals, has changed little in a century. Within this workflow, human salaries are the main cost-driver while data quality is often subject to experimenter variation, and has been difficult to standardize (MacArthur Clark 2017). The result is low statistical power and no technical replicates or comparison with competitor compounds. What is needed is an unbiased, cost effective, repeatable, extensible and scalable paradigm that is built to learn the mapping from animal data to human outcomes and can detect possible failures and blockbuster potential.

Alternatives to *in vivo* rodent pharmacology do not fill the gap despite strong efforts and incentives (FDA 2025). *In vitro* systems such as organoids and organ-on-a-chip have advanced rapidly and now recapitulate tissue-level physiology with increasing fidelity (Low et al. 2020). But a cell culture, however sophisticated, cannot report on the integrated physiology of a behaving animal. Appetite, nausea, sedation, motor coordination, and the emergent behavioral signatures that a drug produces in a living body cannot be measured without movement and an intact nervous system. Larger mammals capture that complexity but do not scale in cost or throughput and raise ethical questions. Moving to humans sooner is not an adequate answer when testing a new compound. Rodents occupy the unique position of a mammal that is small enough for scalability while being complex enough to recapitulate the system-level therapeutic effects that *in vitro* systems cannot.

Recently, advances in behavioral phenotyping have transformed the scale and fidelity with which animal behavior can be measured. Continuous home-cage monitoring has replaced one-off, handling-dependent behavioral assays with uninterrupted measurement across complete circadian cycles (Jhuang et al. 2010; Voikar & Gaburro 2020; Kahnau et al. 2023). In parallel, AI has transformed animal behavior from a set of hand-scored categories into a dense, quantitative signal through markerless pose estimation, action classification, and unsupervised behavioral decomposition (Mathis et al. 2018; Pereira et al. 2022; Weinreb et al. 2024; Wiltschko et al. 2015; Bohnslav et al. 2021). What has been missing is a way to connect that signal directly to the outcomes that matter in the clinic and at a scale sufficient to learn cross-species mappings.

Here we describe a phenotyping platform that records behavior continuously across a fleet of home cages, extracts thousands of behavioral features per hour with an ensemble of computer-vision models, and maps features onto human clinical endpoints by training directly on curated clinical trial data. From a single 24 hour rodent recording, the platform predicts human gastrointestinal adverse-event rates, cardiac and neuropsychiatric toxicity, and long-term clinical weight loss. Despite being 20 times faster than standard multi-week *in vivo* experiments, predictions of human outcomes exceed those obtained from standard long-duration rodent experiments. Because the mapping is unbiased and learned rather than assumed, the same platform extends to further endpoints as clinical data accrue, repositioning the rodent as a directly calibrated model of the human clinic.

## Results

### A scalable *in vivo* platform for rapid experimentation and quality control

The platform begins with a purpose-built phenotype acquisition device (PAD) designed for continuous, unattended behavioral recording at low unit cost (Fig. 1a–c). Each cage integrates an overhead camera, quick-release food hoppers and water bottle, a running wheel, environmental (CO_2_/temperature/humidity) sensing, and a Raspberry Pi. A quick-disconnect trapdoor and magnetic latches increase speed and decrease the effort of routine husbandry (Fig. 1a). The PADs are inexpensive and may be operated in dense parallel arrays (Fig. 1b and c).

**Figure 1.**
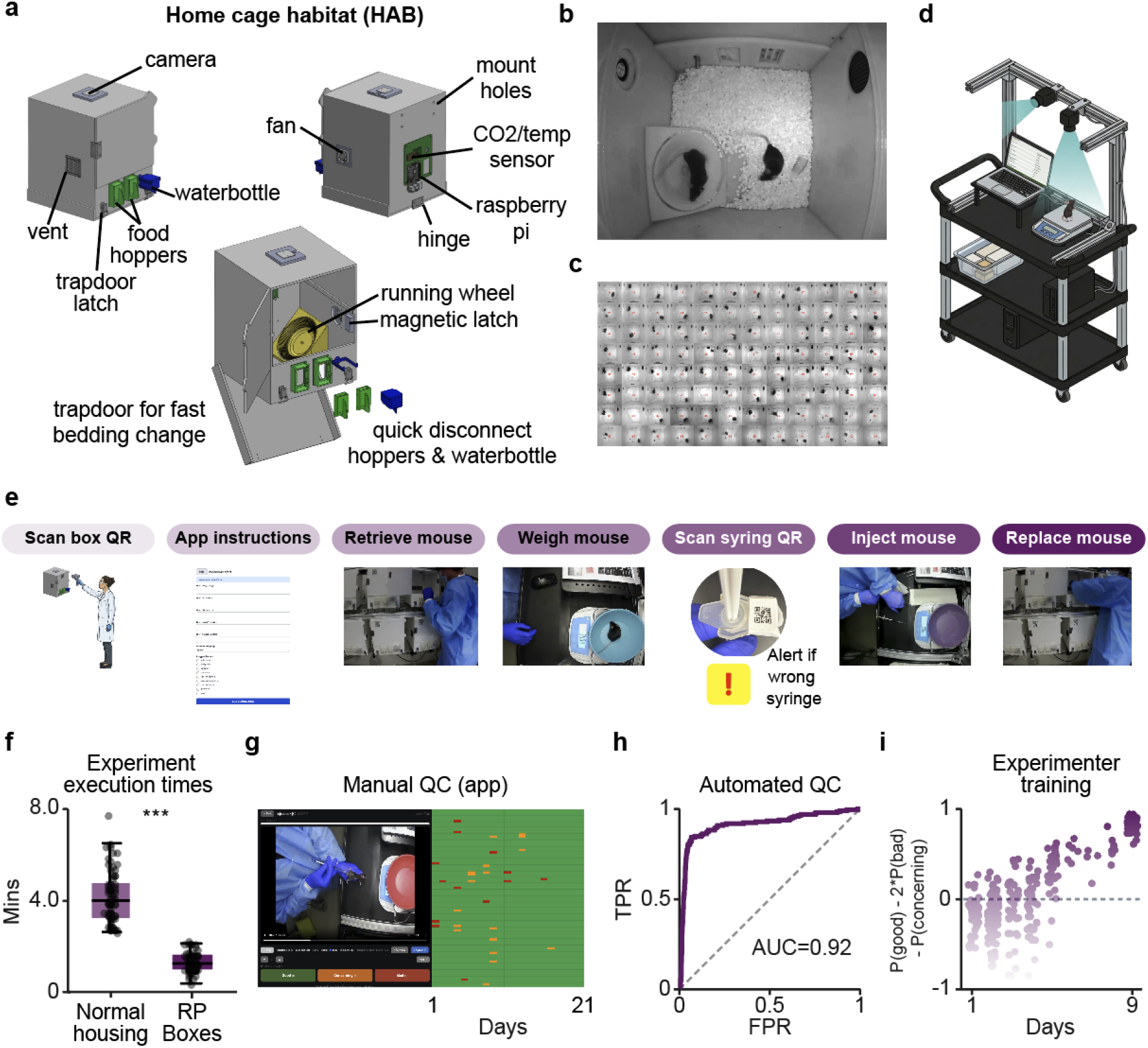
A novel system for quality controlled *in vivo* data at scale. a) Design of an inexpensive and scalable home cage phenotyping acquisition device (PAD) for phenotyping mouse behavior. b) View from IR camera into PAD. c) View from 96 PADs operating in parallel. d) Custom measurement cart used to log food, water, and body weight data while recording animal chain of custody and drug administration. e) Diagram of experimental workflow and tracking. f) Quantification of weighing plus injection times using normal housing (Innovive IVC housing, n=77) or PAD (n=80) (***p<0.0001, Wilcoxon rank-sum). g) Quality Control App used to manually quantify injection quality (left) and example of scored performance over a 3 week experiment. Green blocks: good injection, yellow blocks: concerning, red blocks: bad or wrong route of administration (n=3,360 injections, 99.0% rated “good”). h) Performance of a video classification model trained on 2,577 manually scored injection examples to assess injection quality. g) 9 day example period tracking the performance of a new technician. Drug administration videos and quality scores were used to improve technique of a new technician.

A central source of error in animal experiments is not the measurement itself but the handling around it: dosing the wrong animal, mislabeling a cage, or administering an injection inconsistently. We addressed this with a custom measurement cart and QR code-gated workflow (Fig. 1d,e). The cart contains a scale, 2 cameras, barcode scanner, computer and QR-coded syringes for recording food, water, and body-weight data while simultaneously logging animal chain-of-custody and drug administration. Custom software walks the experimenter through each step in a fixed sequence: scan the PAD QR code, follow on-screen instructions, retrieve and weigh the correct mouse, scan the syringe QR code (which raises an alert if the wrong syringe is presented), inject, and replace the animal (Fig. 1e). The PAD design and cart/QR system reduce experiment time by more than 60% relative to conventional housing (Fig. 1f). In addition to increasing throughput and reducing animal stress, the system enables complete randomization across a fleet of PADs, making the experimental workflow more akin to high-throughput cell-based screening than to conventional *in vivo* pharmacology.

To ensure injection quality, we built an app to visually check every injection (Fig. 1g). After enough data had been collected and annotated (>4,000 intraperitoneal injections), we trained a computer-vision model to score injection quality automatically (Fig. 1h). This model can also be used for realtime feedback to train and correct injection technique (Fig. 1i). Because every measurement is attributable to a verified animal, dose, and operator, automatically time-stamped, and preserved as a complete audit trail, the system is engineered to meet the data-integrity and chain-of-custody standards that underpin Good Laboratory Practice (GLP; OECD Principles of GLP; U.S. FDA 21 CFR Part 58).

### Extraction of rich behavioral repertoires for deep phenotyping

Continuous recording across a 96 PAD fleet generates 2,304 hours of video per day. Raw video streams from infrared cameras to a Raspberry Pi and uploads to cloud storage, with experiment metadata tracked separately in a database. Nextflow dispatches processing across elastic compute instances (Fig. 2a).

**Figure 2.**
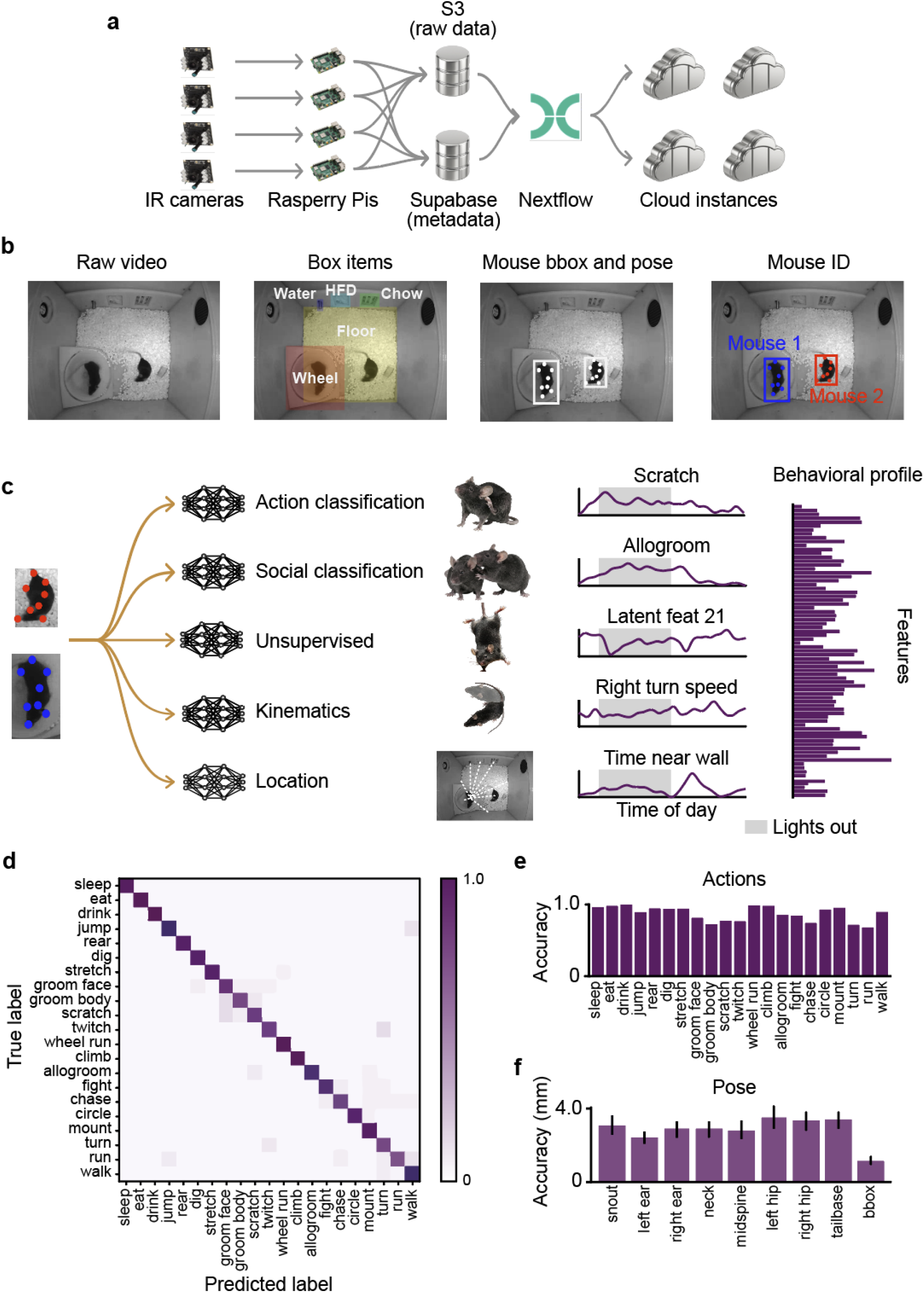
Multi-model extraction of rich behavioral repertoires. a) Diagram of data collection, upload, and processing pipeline. b) Diagram of video data processing steps. Left to right: Raw video image. Objects within the PAD are identified via YOLO model. Mouse pose is identified via YOLO model. Mouse identity is assigned via custom AUC algorithm. c) Far left, middle left: bounding-boxed images and keypoints are sent to downstream models for action and social behavior classification, quantification of unsupervised features, kinematics and location. Middle right: example features over 24 hours (shaded gray area: lights out period). Far right: full feature set creates a behavioral signature for a drug. d) Confusion matrix for action prediction model. Swin 2b model was trained on ∼2.4M labeled actions, and tested on ∼220k balanced labeled actions. e) Accuracies for all behavior classes. f) Accuracy of bounding-boxes and key points using a YOLOv11 model.

We built and trained an ensemble pipeline of computer vision models to extract the full behavioral repertoire of the mouse. The locations of in-PAD items (food hoppers, water spout, floor boundaries, and wheel) are identified with a YOLO (You-Only-Look-Once, Redmon et al. 2016) model; mouse keypoints (nose, ears, neck, spine, hips, tailbase) and bounding-boxes are computed with a second YOLO model, followed by mouse identification (Fig. 2b). We selected YOLO models for their combined bounding-box and keypoint output, inference speed, ubiquity and continued maintenance. Bounding-box-cropped images are fed to an action-classification model that supports both individual and social actions and to an unsupervised model that identifies latent behavioral clusters. Keypoints are used to extract kinematic features (e.g., turn speed) and location (Fig. 2c, left).

Each feature is resolved over the full 24 hour cycle, so that the circadian structure of behavior is preserved. For example, grooming, allogrooming, turning kinematics, and time spent near the cage wall each trace characteristic day–night trajectories (Fig. 2c, second from right). Extracting aggregate values from each feature results in a high-dimensional behavioral signature for each animal, every day (Fig. 2c, right).

Models are continually retrained and updated. For the keypoint and bounding-box YOLO models, we built a custom labeling application and used model-confidence outputs to identify frames requiring manual re-labeling (Supplemental Fig. 1a). Keypoint and bounding-box accuracies are comparable to other published systems (Mathis et al. 2018; Pereira et al. 2022) (Fig. 2f). Distinct actions are identified by a pixel-level video transformer (Swin), trained on over 2.4 million expert-labeled actions and evaluated on approximately 220,000 balanced held-out actions, achieving high per-class accuracy across a repertoire of more than twenty distinct behaviors (Fig. 2d,e). Labels were created using a set of custom apps, one for whole-video labeling of all actions (Supplemental Fig. 1b) and one to rapidly check user- and model-labeled actions using a simple accept/reject two button strategy (Supplemental Fig. 1c).

### Different drugs drive distinct behavioral effects in mice

To create consistent behavioral signatures that can be compared across drugs and connected to clinical data, we developed a standardized experimental paradigm. Animals are maintained on an 8 a.m./8 p.m. light cycle with all dosing performed between 4 and 6 p.m., with behavior recorded continuously (Fig. 3a). Using this protocol, we find that individual compounds shift characteristic subsets of features in distinct directions (Fig. 3b and 3c-n). The antipsychotic olanzapine increases time spent in the middle (open area) of the PAD (Fig. 3c). This increase is not due to locomotor effects as locomotion is decreased (Fig. 3d). Additionally grooming dose-dependently decreases (Fig. 3e). Caffeine is known to have bimodal effects on hyperactivity, with high doses suppressing activity and low doses potentiating it. We indeed see this bimodal effect in turn speeds (Fig. 3f and g) and observe a delay in the onset of wheel running when lights turn off (Fig. 3h), likely due to the depressive effect of high dose caffeine. The CB1 antagonist monlunabant increases grooming (Fig. 3i) and anti-social behaviors like chasing just after injection (Fig. 3j) in a dose-dependent manner. High dose monlunabant also decreased the amount of time mice spent together during lights-on periods when social interaction time is normally elevated, further suggesting antisocial tendencies (Fig. 3k). GLP-1/glucagon co-agonist bamadutide dramatically decreased wheel running, and conversely to monlunabant, increased social interaction time during the lights-out period (Fig. 3l and n). Bamadutide also decreased the amount of time mice spent near walls, suggesting that animals were huddling in the open area of the PAD (Fig. 3m). Dimensionality reduction of all features for this set of drugs reveals that drug exposure drives distinct behavioral changes along similar axes (Fig. 3o), even when baseline values and mouse strains differ (Supplemental Fig. 2a-c).

**Figure 3.**
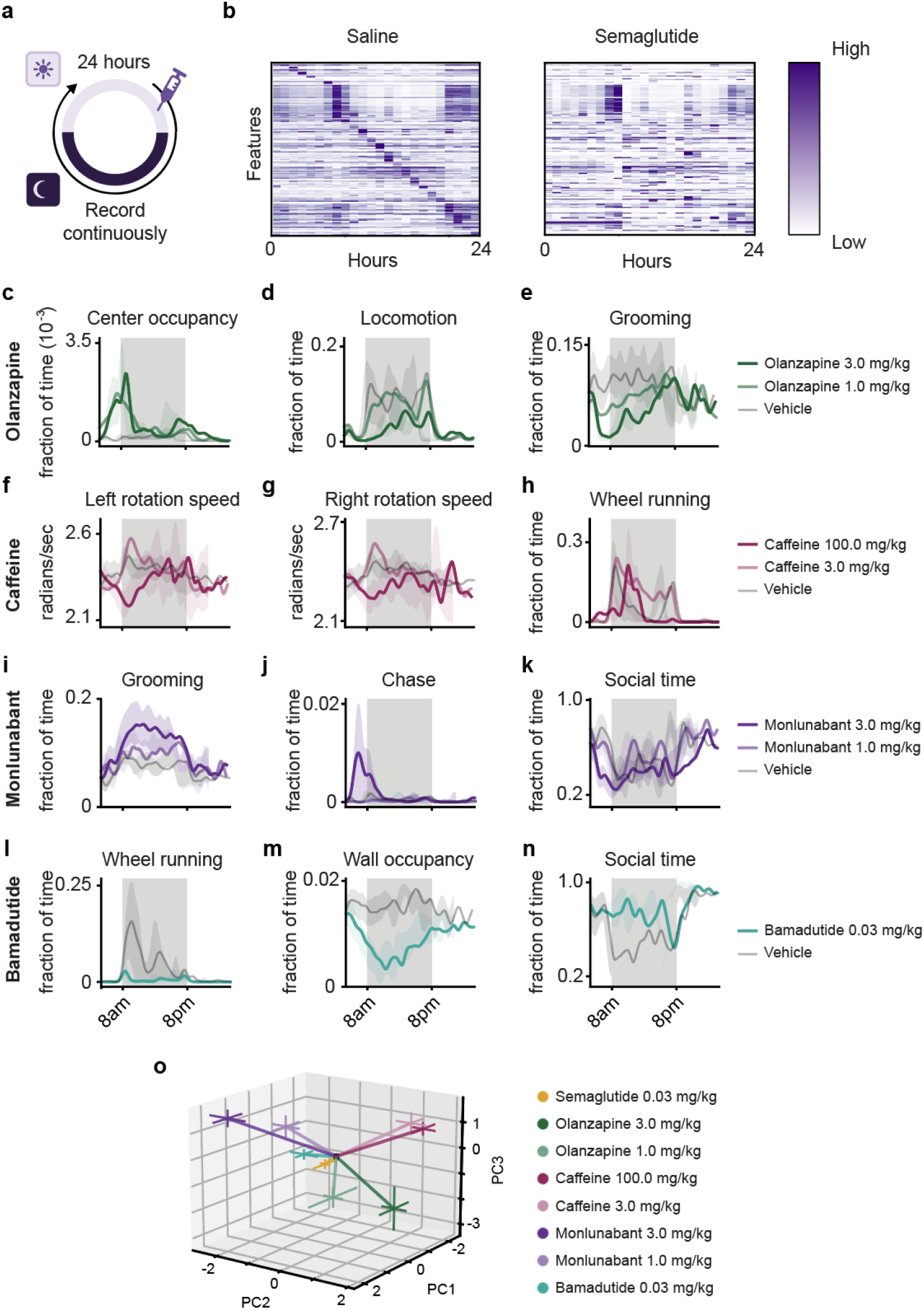
Drugs produce distinct changes in behavior. a) Diagram of 24 hour experimental paradigm. b) Heatmap of features after saline (left) or semaglutide injection (right). c-n) examples of specific features that change relative to controls. Olanzapine (c-e, n=12 PADs per group), caffeine (f-h, n=7-11 PADs per group), monlunabant (i-k, n=7-11 PADs per group), bamadutide (l-n, n=8 per group). o) PCA projection of drugs using all features. Different MOAs occupy different spaces.

### Rodent behavior predicts human clinical outcomes

We hypothesized that behavioral features captured distinct physiological processes arising from drug-induced perturbations that are detected by the nervous system and impact behavior (Fig. 4a). Further, although the underlying effects and mechanisms may be different between mice and humans, the effects of drugs in both species allow an unbiased transform to be computed between them.

**Figure 4.**
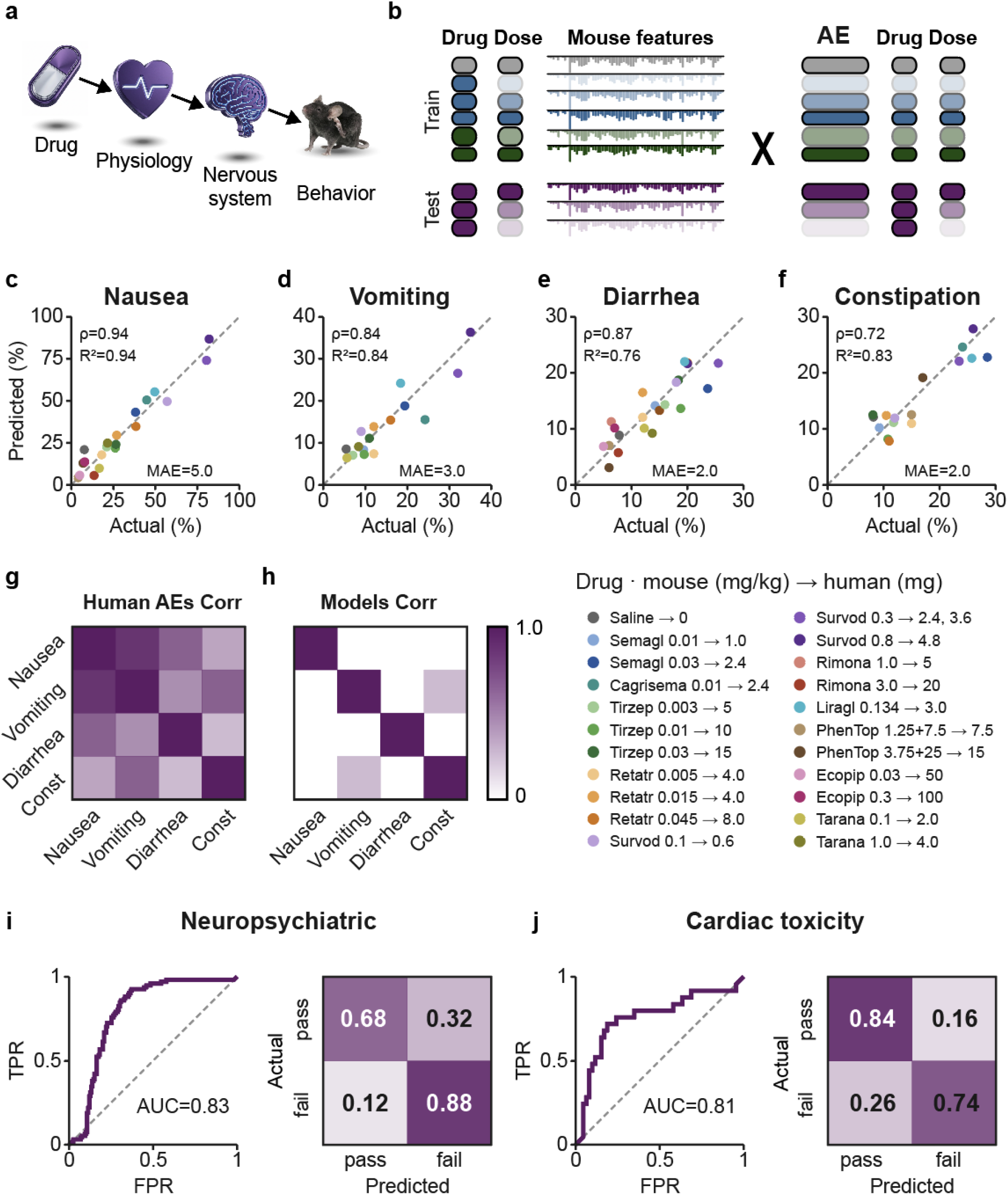
Rodent behaviors predict human outcomes. a) Diagram for how a drug will affect behavior along specific dimensions. b) Diagram of model building. Mouse drug-phenotype fingerprints are paired with human drug-outcome data. c-f). Models trained on rodent behavior to predict rates of (c) nausea (n=22 drug doses, 1404 PADs), (d) vomiting (n=16 drug doses, 649 PADs), (e) diarrhea (n=22 drug doses, 877 PADs) and (f) constipation (n=16 drug doses, 650 PADs) in humans. Labels in lower right. g) Pairwise association among human adverse-event labels from clinical trials. Heatmap values show R^2^ from Pearson correlations of manipulation-level AE rates. All off-diagonal AE pairs were significantly correlated after BenjaminiHochberg FDR correction (R = 0.577-0.933; R^2^ = 0.333-0.870; all q *≤* 4.13e-5), indicating substantial correlation structure among the human AE labels. h) Pairwise association among rodent adverse-event model feature-weight profiles. Heatmap values show R^2^ from Pearson correlations of signed, standardized ElasticNet coefficient vectors. No off-diagonal model-pair correlations were significant after Benjamini-Hochberg FDR correction (all q = 1.0; minimum uncorrected p = 0.287 for vomiting vs constipation), indicating that the fitted rodent feature-weight profiles were not significantly linearly correlated despite correlation among the corresponding human AE labels. (i-j). ROC and confusion matrices (row normalized) for models trained on rodent behavior to predict if a drug will drive neuropsychiatric (i) or cardiac toxicity (j) events in a clinical trial. (neuropsych: F1: 0.720, precision: 0.608, recall: 0.884, accuracy: 0.753 n=) (cardiac: F1: 0.708, precision: 0.680, recall: 0.739, specificity: 0.843, balanced accuracy: 0.791, ROC-AUC: 0.806, n = 52,650 unique mouse:experiment datasets) ranging from 0.70 to 1.00 (median 0.99). Together these results suggest that behavioral phenotyping provides signals for both toxicity and tolerability.

To test this hypothesis, we assembled a database of human outcomes using published clinical trial data from the AACT database (Tasneem et al. 2012), company press releases, and published articles. To structure and normalize this data we built a multi-step AI agent + human pipeline that processes trial records and PDFs to extract arms, doses, schedules, endpoints, adverse events, and eligibility criteria (Supplemental Fig. 3). Human and rodent dosing were paired using an allometric bridge (human dose in mg *≈* rodent dose in mg/kg *×* 3/37 *×* 70 kg *≈* 5.68 *×* rodent dose in mg/kg) and rounded to values found in the literature or clinical trials (Supplemental Fig. 4). For each drug and dose, the resulting mouse behavioral fingerprint is paired with the corresponding human drug–outcome data drawn from clinical trials, and models are trained to map the former onto the latter (Fig. 4b).

To build out the first set of human-outcome predictions, we selected gastrointestinal adverse events in clinical trials for obesity and/or diabetes. This trial set has several advantages: the trials share similar designs, patient populations, and outcome measures, and gastrointestinal adverse events are both a major driver of treatment discontinuation and, in some cases, have halted clinical development. Trained on this data, models built from rodent behavior predict human gastrointestinal adverse-event rates with striking accuracy across a panel of compounds spanning incretin agonists, centrally acting agents, and combination therapy (Fig. 4c–f). Nausea in particular is predicted better than by simply using food-intake reduction as a proxy (Supplemental Fig. 5). Roughly 10 PADs per drug, with 8 or more drugs and 2–3 doses per drug, is sufficient to achieve this level of prediction (Supplemental Fig. 6). The human adverse-event rates are themselves strongly correlated (Fig. 4g), yet the fitted feature-weight profiles for the four endpoints were not significantly correlated (Fig. 4h). Thus, the models do not rely on a single shared set of behavioral features, suggesting they capture distinct, outcome-specific signatures rather than a generalized tolerability axis.

We next tested if rodent behavioral data could predict clinical failures for neuropsychiatric side effects (Fig. 4i). Models were able to accurately predict failed drugs including rimonabant, taranabant, and ecopipam (still in development for Tourette’s, but discontinued for obesity; Astrup et al. 2007). The top features were increased grooming and scratching while positive social interaction decreased — all features associated with neuropsychiatric indications in rodent models but often missed without 24 hour observation (Kalueff et al. 2015). Interestingly, setmelanotide and phentermine/topiramate were predicted to have a higher probability of psychiatric failure than other drugs in the dataset. Although they are approved, both carry label warnings for neuropsychiatric side effects, demonstrating a gradient in predictions.

We hypothesized that more subtle toxicity signals would also surface in high dimensional mouse behavior. To this end, we tested a separate panel of drugs that had failed clinical trials for cardiac toxicity (Fig. 4j). The panel consisted of 74 drug-dose conditions, including 23 cardiac-toxicity positives (18 cardiac and 5 hERG-associated conditions) and 51 safe controls. Performance was particularly strong for hERG-associated liabilities (Recanatini et al. 2004): all five hERG-positive conditions were flagged as positive in held-out predictions, with predicted failure probabilities

### The platform accelerates development and improves translation

Standard rodent weight loss studies are two to three weeks long and track body weight daily. We compiled data from this paradigm across published literature to determine how well this protocol predicts human weight loss (Fig. 5a), and compared the results to predictions generated from our platform’s 24 hour behavioral data (Fig. 5b). Predictions based on single-day behavioral features correlated more strongly with human data (Fig. 5c) and reduced the median absolute error (MAE) by approximately 70% (Fig. 5d). A day of behavioral data thus recovers the human-relevant signal more accurately than three weeks of conventional weight tracking.

**Figure 5.**
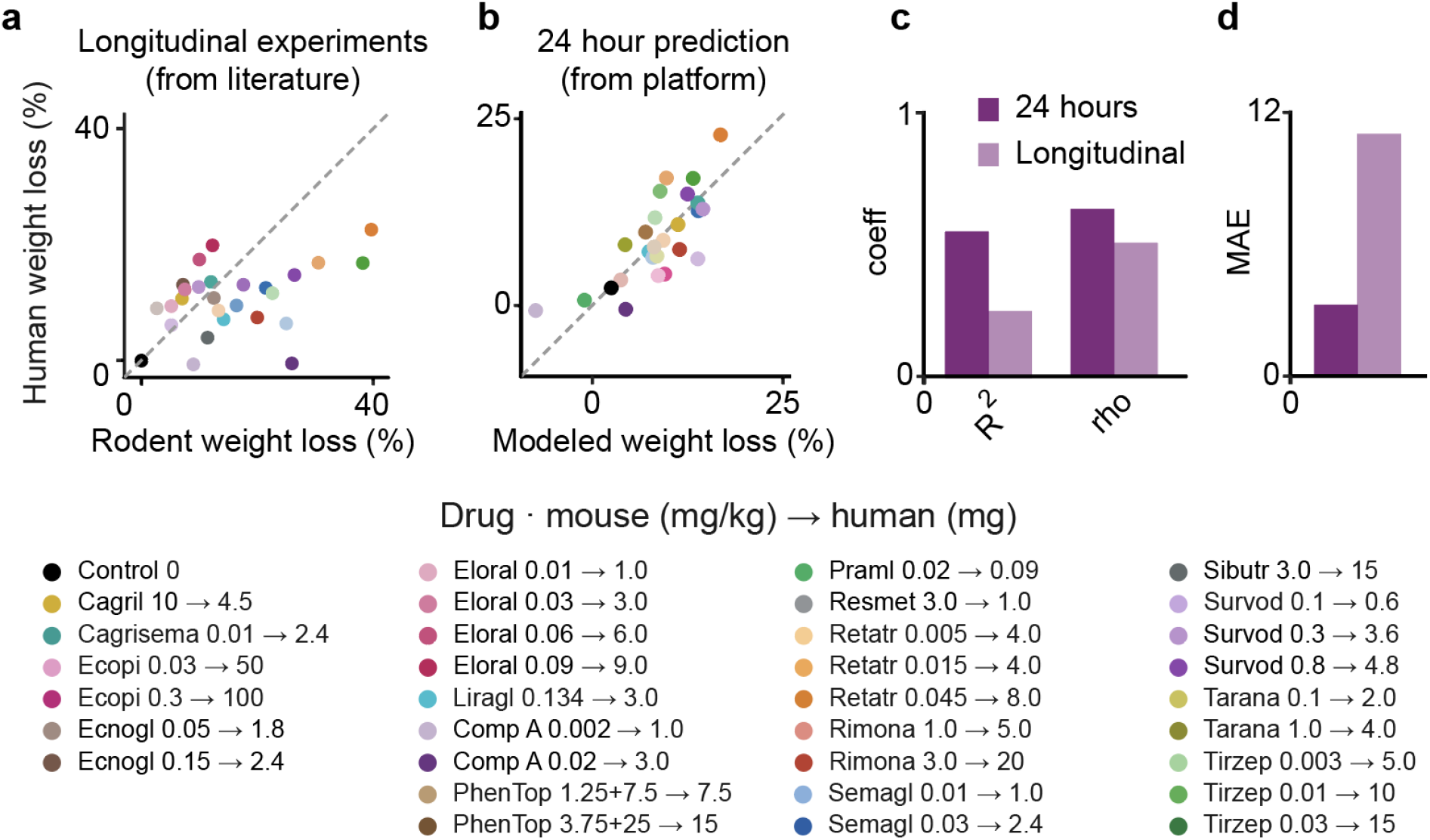
24 hours of behavioral data outperforms longitudinal experiments for predicting human weight loss. a) Published rodent weight loss at study endpoints (2-3 weeks) vs reported human clinical weight loss at weeks 40 through 80. b) Predicted weight loss from 24 hour behavioral data vs reported human clinical weight loss (same as a). c-d) Comparison of human-outcome correlation (Pearson r2 Spearman rho, c) and error (MAE, d) for the overlapping drugs/doses across published longitudinal experiments vs predictions from 24 hours of behavioral data.

This combination of speed and fidelity reframes what a preclinical experiment can do. Mapping the average clinical gastrointestinal adverse-event (GI-AE) burden against clinical weight loss for four generations of best-in-class weight-loss therapies (liraglutide Pi-Sunyer et al. 2015, semaglutide Wilding et al. 2021, tirzepatide Jastreboff et al. 2022, and retatrutide Jastreboff et al. 2023) defines an efficacy–tolerability landscape in which successive generations occupy a more favorable position, approaching the optimal lower-left corner characterized by greater efficacy and reduced GI-AE burden (Fig. 6a). The platform is able to recapitulate this efficacy–tolerability landscape and evaluate new therapeutics on a 24 hour timescale (Fig. 5b). To demonstrate this, we used a computational algorithm to generate sets of compounds that were predicted to surpass semaglutide or retatrutide in efficacy and tolerability across seven 24 hour iterations. Results were fed back into the algorithm to generate a modified set of compounds to test the next day. Over this iterative process efficacy-tolerability predictions moved down and left, passing retatrutide after day 4 (Fig. 5c). These results demonstrate that the platform can be used for multiparameter optimization on a daily timescale.

**Figure 6.**
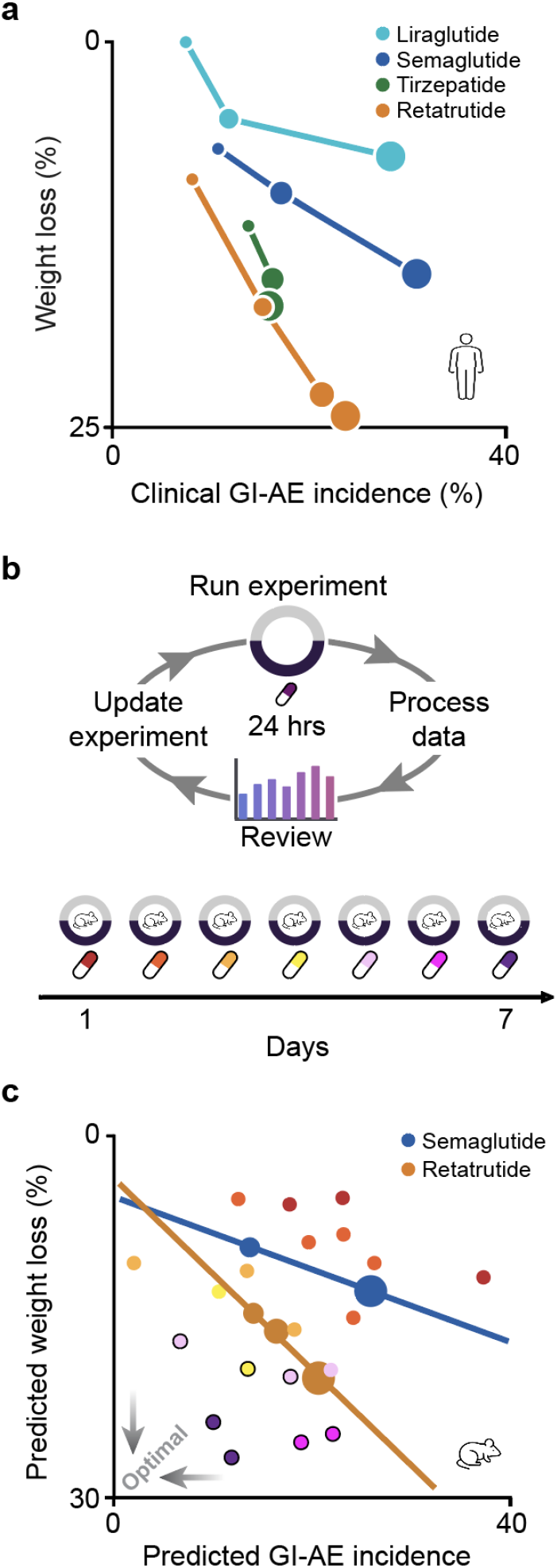
Daily feedback loops promote rapid discovery of superior candidate therapies. a) Clinical results for average adverse events reported vs weight lost, across four generations of best-in-class weight-loss therapies. Superior therapies trend towards the bottom left, showing better efficacy:tolerability ratios relative to prior therapies. b) Schematic of Olio’s 24 hour data collection, analysis, and planning system (top) and timeline for a week of Olio experiments testing candidate weight-loss compounds (bottom). c) Results from 1 week of iterative experimentation, color coded by date. On each day, 2-3 different drug/dose candidates were tested, and compared to internal semaglutide and retatrutide data.

## Discussion

### Behavior reflects physiology

The framework is founded on a mechanistic chain linking molecular target engagement to physiology and, ultimately, behavior. A drug perturbs molecular pathways and, through them, the body’s physiology. Physiological changes are continuously sensed by the nervous system, which acts like a high-bandwidth antenna. Virtually any shift in internal state, whether metabolic, cardiovascular, gastrointestinal, immune, is registered centrally, and any change in the nervous system’s state will ultimately be expressed as a change in behavior. Multiple nonlinear processes including pharmacokinetics, biodistribution, target occupancy, and biased agonism separate a drug’s nominal target from its physiological and behavioral effects. Consequently, even drugs that share a target, such as the GLP-1R agonists liraglutide and semaglutide, produce distinct, structured behavioral signatures in our system. Behavior is therefore not a crude proxy for a single axis of illness but a high-dimensional readout of systemic pharmacology.

The premise that physiological state is written into behavior is well established. Across species, infection and inflammation produce a coordinated, centrally organized set of behavioral changes (reduced activity, altered feeding, social withdrawal) that are not incidental but an adaptive program orchestrated by the nervous system (Hart 1988; Dantzer et al. 2008). The same logic extends to organ-specific pathology rarely thought of as “behavioral”: heart failure and cardiac dysfunction manifest as exercise intolerance and fatigue resulting in measurable reductions in physical activity (Del Buono et al. 2019). In older adults a urinary tract infection commonly presents first as confusion or delirium, rather than as a classical localized symptom (Dutta et al. 2022). If one could record and quantify every aspect of physiology and behavior an organism performed, all pharmacological and disease perturbations would leave a detectable trace.

Human physiology is already being read this way, at scale, with far coarser instruments than those presented here. Consumer wearables using little more than an accelerometer and an optical heartrate sensor recover sleep architecture with incredible accuracy (Kanady et al. 2020), can detect deviations from personal physiological baselines that precede overt illness (Li et al. 2017), and flag previously undiagnosed atrial fibrillation (Perez et al. 2019). Continuous, high-resolution video of an animal’s entire behavioral repertoire should carry an even richer picture of its internal state.

### Animal studies are biased

In a conventional rodent behavioral assay the experimenter first decides which behavior ought to correspond to a human clinical state, then measures that one behavior and interprets it as a stand-in for the state (Shemesh & Chen 2023). This design embeds two assumptions that are rarely tested directly and have resulted in decades of poor clinical translation. The first is that the chosen readout actually maps onto the human condition (Nestler & Hyman 2010; Belzung & Lemoine 2011; Wong et al. 2010). The second is that an animal’s internal state can be determined from simple pre-specified readouts (Anderson & Adolphs 2014). For example, the forced-swim test is routinely read as “behavioral despair,” but its central measure, immobility, could as easily be driven by energy demands as it is a depression-like affective state (Cryan et al. 2005; Molendijk & de Kloet 2019). Even a measure as seemingly straightforward as weight loss suffers from the one-dimensional readout. Weight loss can be caused by many changes including dehydration, or muscle, fat and bone-density loss. By including all behavioral features we obtain more accurate human weight prediction than using mouse weight change alone (Fig. 5d), demonstrating the power of unbiased, high-dimensional representations when predicting human outcomes from animals.

We make no a priori assumptions about which behavior matters or about what the animal is experiencing. By measuring the full behavioral repertoire without privileging any feature, and building models that predict human clinical data directly, we let the models empirically select features that carry outcome-relevant information. An interpretation of the mouse’s internal state is not required to make predictions, and the model feature weights can be interrogated for interpretation — there is no black box.

### *In vivo* studies as high-throughput screens

Rather than replacing conventional confirmatory animal studies, the platform occupies a different role in the discovery pipeline. It functions more like a high-throughput cell-based screen than a traditional *in vivo* experiment. Instead of testing a few doses against a control across several endpoints, it enables parallel comparisons of many compounds, technical replicates, dose–response profiling, and yields quantitatively comparable data for ranking and selection. Therefore the natural place to deploy it is when a program has multiple plausible candidates and no principled way to rank them. All can be run in parallel, along with competitors and scored on a multidimensional therapeutic index. For example, the platform can answer whether a candidate delivers meaningful weight loss with fewer side effects. Liraglutide and semaglutide illustrate this. Both target the same receptor but have very different effects on weight loss and gastrointestinal adverse event rates. Because experimental cycles are 24 hours, candidates can be iterated as quickly as they can be synthesized, giving *in vivo* discovery the cadence of high-throughput screening rather than of multi-week animal studies. This is especially advantageous in the era of agentic experimental and molecule design.

### Ethical considerations

The platform changes the ethical calculus of animal research by advancing the 3Rs—replacement, reduction, and refinement (MacArthur Clark 2017). By making rodent behavior directly predictive of human outcomes, it may replace some studies in dogs and non-human primates with studies in a lower-sentience species (replacement). Each recording yields thousands of behavioral features spanning many potential endpoints, allowing fewer animals to answer more questions. Longitudinal monitoring and within-PAD modeling further increase statistical resolution, while enabling each mouse to be studied across multiple compounds (reduction). Finally, passive home-cage monitoring avoids the handling and novel-environment stress associated with conventional assays (refinement; Voikar & Gaburro 2020; Kahnau et al. 2023).

The infrastructure also supports welfare and compliance directly. Automated husbandry alerting (flagging weight loss, inactivity, or other deviations in real time) allows problems to be caught earlier and more consistently than intermittent human checks. More broadly, computer-vision monitoring can substitute for the continuous human observation that regulated studies require. Because every action is attributed, time-stamped, and archived, the resulting records meet the data-integrity standards that underpin GLP by construction. In principle this makes it feasible for every experiment to be GLP-grade by default, rather than reserving that rigor for a small number of costly pivotal studies, further reducing the number of experiments required for drug development. Given this, and the speed and efficiency of the platform, we would argue that all rodent studies would benefit from being run on such a platform. The marginal cost of capturing the full behavioral record is negligible, while the information forgone by not capturing it is large.

### Limitations

While the method described herein is flexible and extensible, limitations still bound the approach. First, the predictions are only as good as the human data they are trained on; heterogeneous trial designs, inconsistent adverse event documentation, and sparse coverage of some endpoints set a ceiling on accuracy that no amount of rodent data can overcome. Improving the quality and standardization of the clinical training corpus is as important as scaling animal data (Ioannidis 2009; Golder et al. 2016). Second, the approach depends on cross-species conservation of the relevant biology: where a drug’s target is not functional in the mouse, behavioral readouts cannot carry information about it. A clear example is the non-conservation of a receptor. Some nonpeptide GLP-1R agonists exhibit strong species selectivity because of receptor-sequence differences, motivating use of humanized GLP-1R models (Zhang et al. 2020; Jun et al. 2014). Importantly, though, the platform tolerates quantitative species differences as long as the drug still perturbs mouse behavior, because the learned mapping accounts for species-specific scaling and pathway differences rather than assuming mechanistic identity. For example, amylin pharmacology differs markedly between mouse and human (Hay et al. 2015; Duffy et al. 2018), yet the platform nonetheless predicts and correctly ranks cagrilintide and cagrilintide–semaglutide weight loss and adverse-event rates. The boundary, then, is not species differences per se but targets that are behaviorally silent in the mouse. However, care is warranted whenever a candidate’s target or downstream pathway physiology is poorly conserved.

### Extensibility

The mouse is, for now, uniquely suited for the 24 hour experiment. Metabolic rate scales inversely with body size across mammals which results in time-compressed physiological changes in mice relative to larger species (West et al. 1997). A physiological process that would take days to play out in a larger species can surface within a single day of recording in the mouse. This speed comes with a well-known cost: mice do not share the pharmacokinetic profiles of humans, which are better approximated in rats and larger mammals. Interspecies differences in drug clearance and exposure are substantial enough that they are routinely corrected for by allometric scaling (Mahmood et al. 2006; Mahmood 2007). The mouse thus trades pharmacokinetic fidelity for temporal compression. This is acceptable because the platform learns the behavior-to-outcome mapping directly rather than relying on matched exposure. Long-term experiments are, of course, still possible in the platform. However, the predictive validity of an assay must be demonstrably increased to justify daily dosing of animals for many weeks.

Nothing about the underlying system is specific to the mouse. The imaging, pose-estimation, action-classification, and feature-extraction pipeline is species-agnostic in principle, and comparable computer-vision methods have already tracked posture and behavior in macaques (Bala et al. 2020). Porting the platform to larger species is therefore a matter of hardware design, retraining the vision models on the new animal and mapping non-human to human dosing to connect clinical data.

## Conclusion

Herein we introduced a novel platform that directly links animal data to human outcomes in an extensible and unbiased framework. We expect the repertoire of predictions to grow substantially, first through additional endpoints in mice and, ultimately, through adaptation to other species, expanding the utility and predictive validity of animal models across drug discovery and development.

## Methods

### Animals and husbandry

Protocols were approved by an external Institutional Animal Care and Use Committee. Male and female C57BL/6 mice were purchased from Jackson Laboratories and Taconic Biosciences at age 8 to 12 weeks. Mice were habituated in PADs equipped with a running wheel in a 12/12 light cycle for 10 days and fed ad libitum water, high fat diet (HFD, Research Diets D12492i) and regular chow (Labdiet Picolab 5053) in separate hoppers. Each PAD housed 2 mice of the same sex. Behavior was observed over at least 7 days before mice entered experiments. In cases of abnormal behavior or fighting, pairs of mice would be replaced.

### Behavioral feature extraction

Recordings were saved to AWS S3 with metadata about the experiment saved to Supabase. PAD items (wheel, hoppers, water spout, floor edges) are extracted from one initial frame at the beginning of a recording. On a frame-by-frame basis the mice are bounding-boxed and key points identified via YOLO model. A custom algorithm identifies each mouse from the prior frame so bounding-box jumps are minimized. Cropped frames of each mouse are processed by a Swin 2b vision transformer. Key points are further processed into kinematic and location features. Action and social action frequencies, locations, and kinematics are binned in 5 minute or 1 hour bin files for model training. Data presented in Figure 3c-n is from 5 minute bins smoothed with a 30 minute Gaussian window. PCA in Figure 3o is from 5 minute bins and data normalized within the PAD.

### Clinical-trial data curation

AACT reports and pdfs from clinical trials were processed first by a deterministic data extractor (https://aact.ctti-clinicaltrials.org/) and an agentic extractor (pdfs). Data was structured by trial, which was then broken into trial arms and study criteria. Arms were then broken into drugs and doses, schedules, endpoints, adverse events and baseline data. Deterministic normalization was used to categorize terms. After the data was fully curated, every value was verified by a human before being used in prediction. We selected U.S.-based obesity and diabetes clinical trials, focusing on Phase II and later, with durations of 40–80 weeks. We included studies with participants aged 18–60, ensuring a mix of male and female subjects, and did not exclude trials based on dose-ramping protocols. If multiple trials were available for a single drug (such as the STEP trials for semaglutide) we averaged the adverse events and endpoints across them, alongside placebo data.

### Historical rodent weight loss data

Published data in Figure 5 was scraped from papers published on Pubmed for longitudinal (2 or 3 week) weight loss data for drugs listed in Figure 5. If the mean or median weight loss percent was not listed in the text, we made a best estimate using the figure for the ending weight for a given cohort.

### Predicting human gastrointestinal adverse events and weight loss

Gastrointestinal endpoints (nausea, vomiting, diarrhea, constipation) and weight loss were modeled by regression against curated human adverse-event rates. Within PAD data was normalized to non-injection, washout or habituation days preceding the experimental day. Models were standardized elastic-net regressions; regularization was selected by internal cross-validation over a range of L1/L2 mixing values with a fixed random seed. Generalization was assessed by leave-one-out cross-validation, refitting scaling and model in each fold, and performance was reported from outof-fold predictions using Pearson correlation, its square (R^2^), and Spearman correlation. After preprocessing and quality control, endpoint datasets comprised 22 conditions for nausea (from 1,404 enclosure-level observations), 16 for vomiting (from 649), 22 for diarrhea (from 877), and 16 for constipation (from 650).

### Neuropsychiatric and cardiac toxicity

Cardiac-toxicity and neuropsychiatric-failure liabilities were each modeled as binary classification against curated human labels, sharing a common design: behavioral preprocessing as above, evaluation by held-out cross-validation in which all observations sharing a label unit were held out together, and reporting from out-of-fold predictions by ROC AUC with F1, precision, recall, accuracy, and confusion matrix. The two endpoints differ in their unit of prediction and label source, as follows. For cardiac toxicity, positive conditions were compounds annotated with cardiac-toxicity or hERG-channel liability; controls were vehicle or drug conditions annotated as safe with no known toxicity label, and conditions with missing or non-cardiac annotations were excluded from the primary analysis. The positive set comprised 26 conditions before feature and quality-control filtering and 23 in the primary analysis (18 cardiac and 5 hERG); representative positives were well-characterized cardiotoxic and hERG-active agents such as cisapride, terfenadine, rofecoxib, sibutramine, dofetilide, and E-4031. The control set comprised 57 conditions before filtering and 51 in the primary analysis, spanning vehicle controls and diverse pharmacological comparators across metabolic, central-nervous-system, analgesic/anti-inflammatory, and cardiovascular classes. Several classifier families were compared against a baseline (regularized logistic regression, random forest, gradient-boosted trees); the selected model was a gradient-boosted-tree classifier with treatment-condition-balanced weighting, chosen by held-out F1 and precision–recall AUC, with confidence intervals from bootstrap resampling of held-out condition predictions.

For neuropsychiatric failure, weight-loss compounds that failed clinically for neuropsychiatric reasons were distinguished from those without known neuropsychiatric failure, using literature-derived labels and the highest available dose per drug. The L2-regularized logistic regression with class-balanced weighting was evaluated by leave-one-drug-out cross-validation; the regularization strength was tuned by nested leave-one-drug-out selection on the training drugs within each outer fold, and enclosure-level probabilities were averaged within each held-out drug to yield drug-level predictions. Signed model coefficients were used for feature-importance summaries.

## Supplemental Figures

**Supplemental Figure 1.**
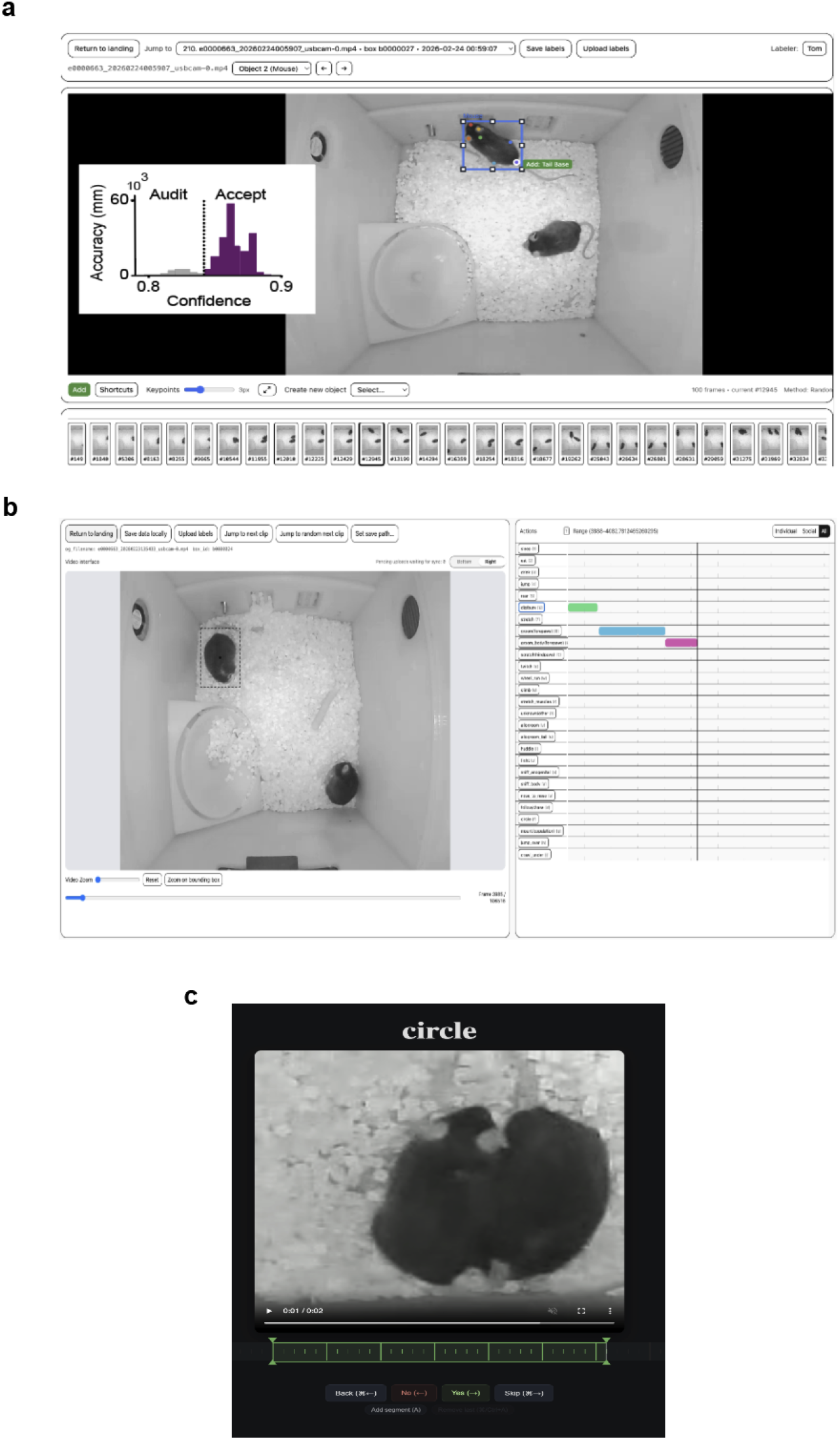
Apps used for data labeling. a) App for labeling bounding-boxes and keypoints, and histogram of point confidences used for human-in-the-loop iteration. b) App for labeling behavioral classification. c) App to rapidly QC labeled or inferred actions using a strategy similar to online dating apps .

**Supplemental Figure 2.**
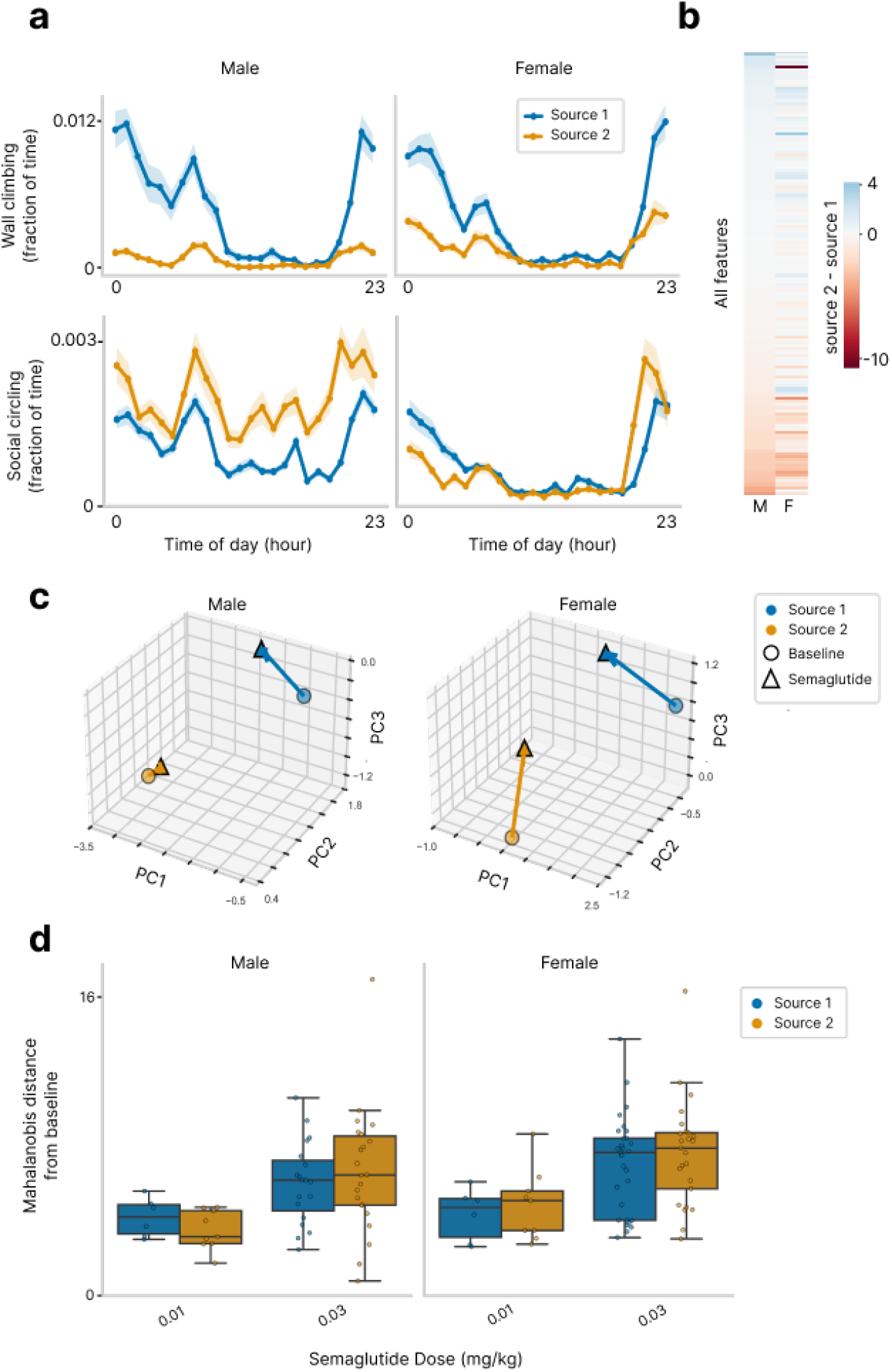
Mouse source differences and drug response similarities. Olio platform detects differences in mouse strains, but maintains drug sensitivity. a) Comparison of two example behavioral features for mice sourced from two different vendors. Mice were age-matched and recorded on the same days. b) Comparison of Source 2 - Source 1 aggregate feature values for male (left) and female (right) mice. c) Latent (PCA) feature representation of mice from both sources under baseline conditions and semaglutide dosing. d) Mahalanobis distance between baseline and semaglutide conditions across doses. No significant differences between sources were observed (Mann-Whitney U).

**Supplemental figure 3.**
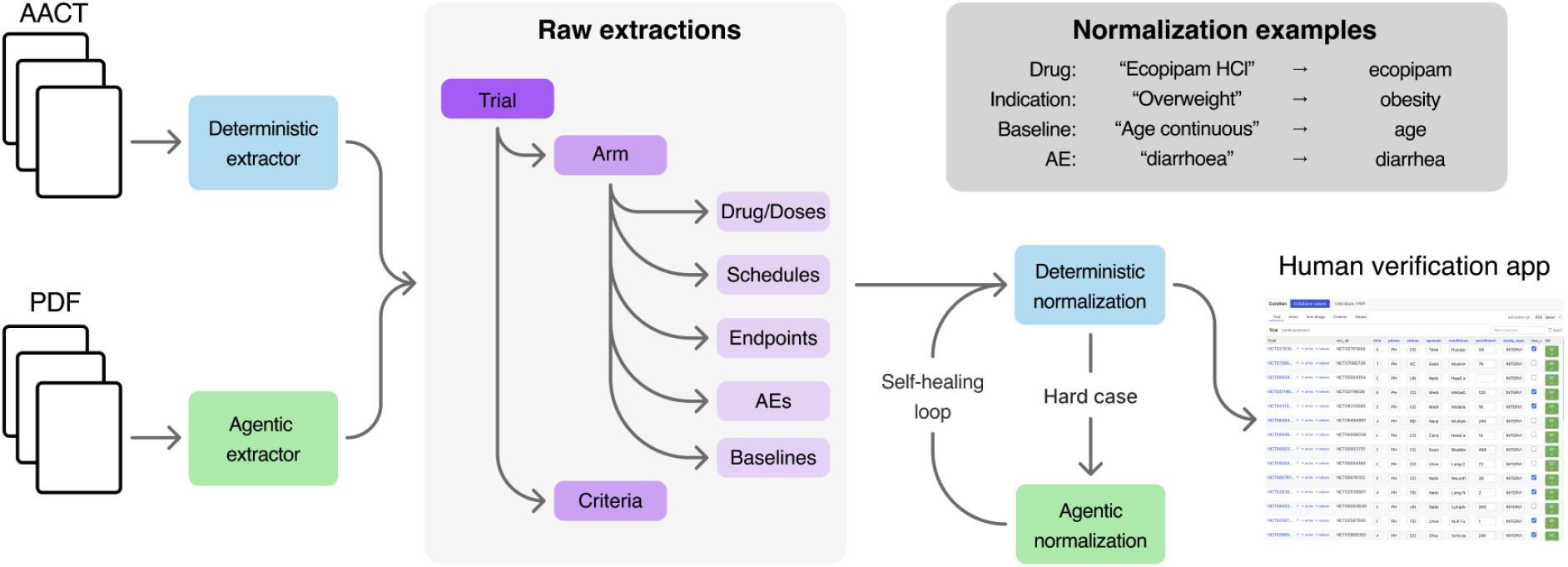
Agentic and human clinical data curation. Schematic of workflow. AACT reports and pdfs from clinical trials are processed first by a deterministic data extractor (AACT) and an agentic extractor (pdfs). Data is structured by trial, which is then broken into trial arms and study criteria. Arms are then broken into drugs and doses, schedules, endpoints, adverse events and baseline data. Deterministic normalization is used to categorize like terms with an off-ramp to an agentic normalization for hard cases that 1) creates a new normalization and 2) patches the deterministic normalization for future runs. After the data is fully curated, every value is verified by a human before being used in prediction. Verified data is also used for further training deterministic and agentic extractors and normalizers.

**Supplemental Figure 4.**
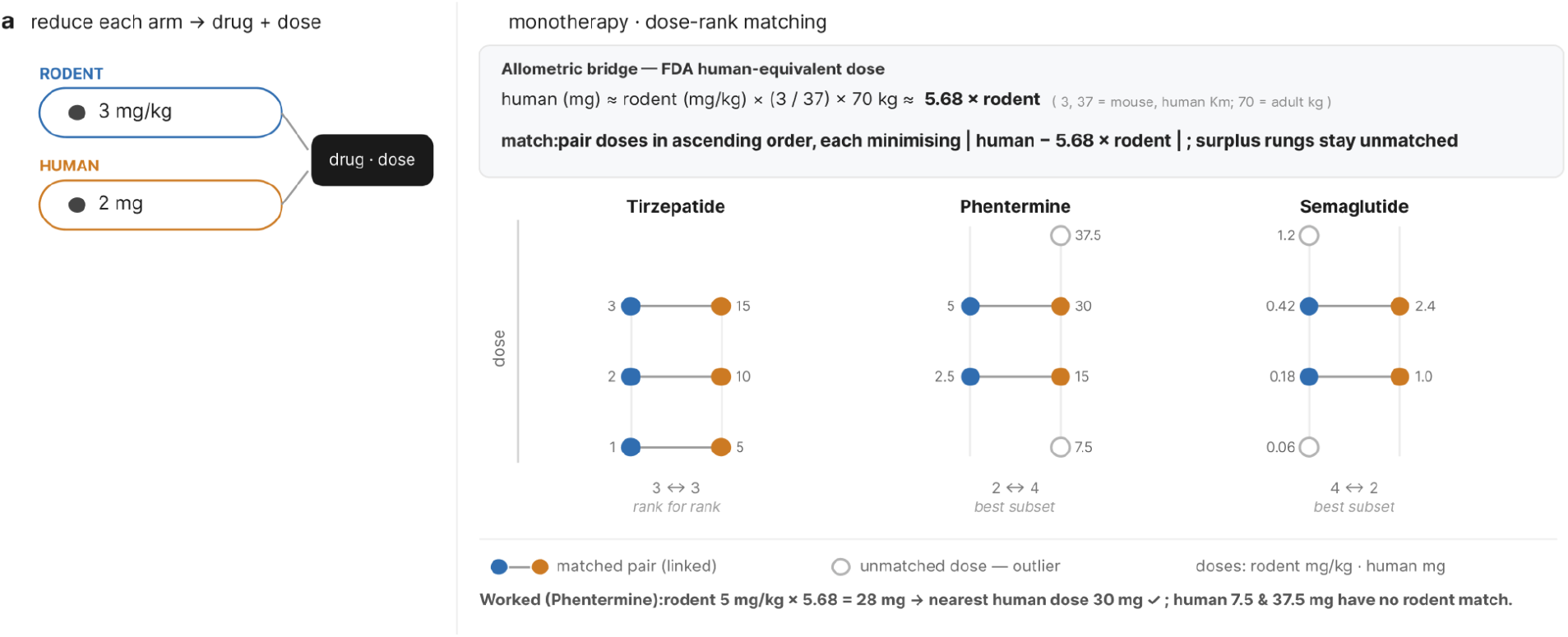
Mapping human and mouse drug doses.

**Supplemental figure 5.**
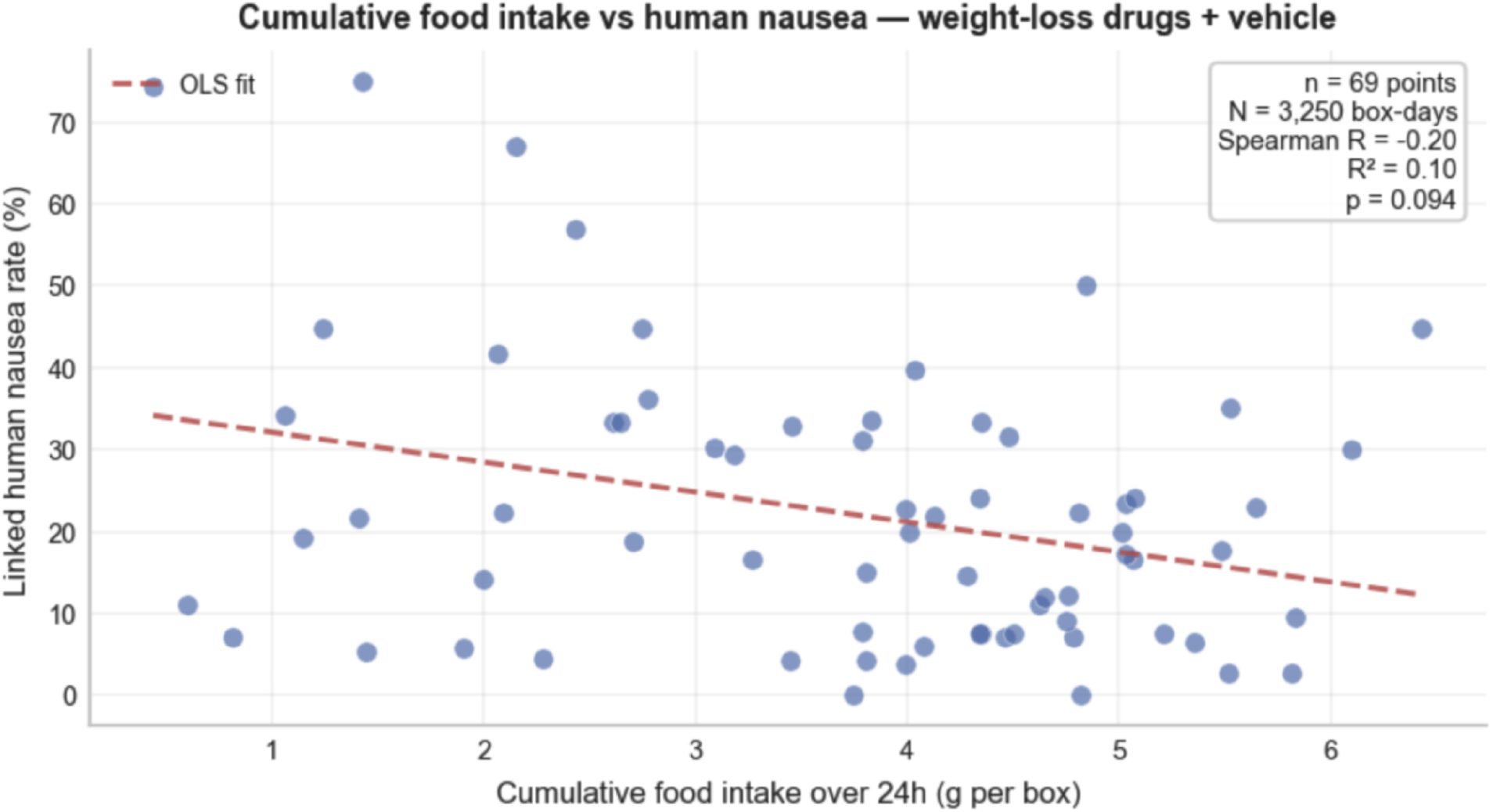
Food intake suppression is not a reliable predictor of human nausea rates.

**Supplemental figure 6.**
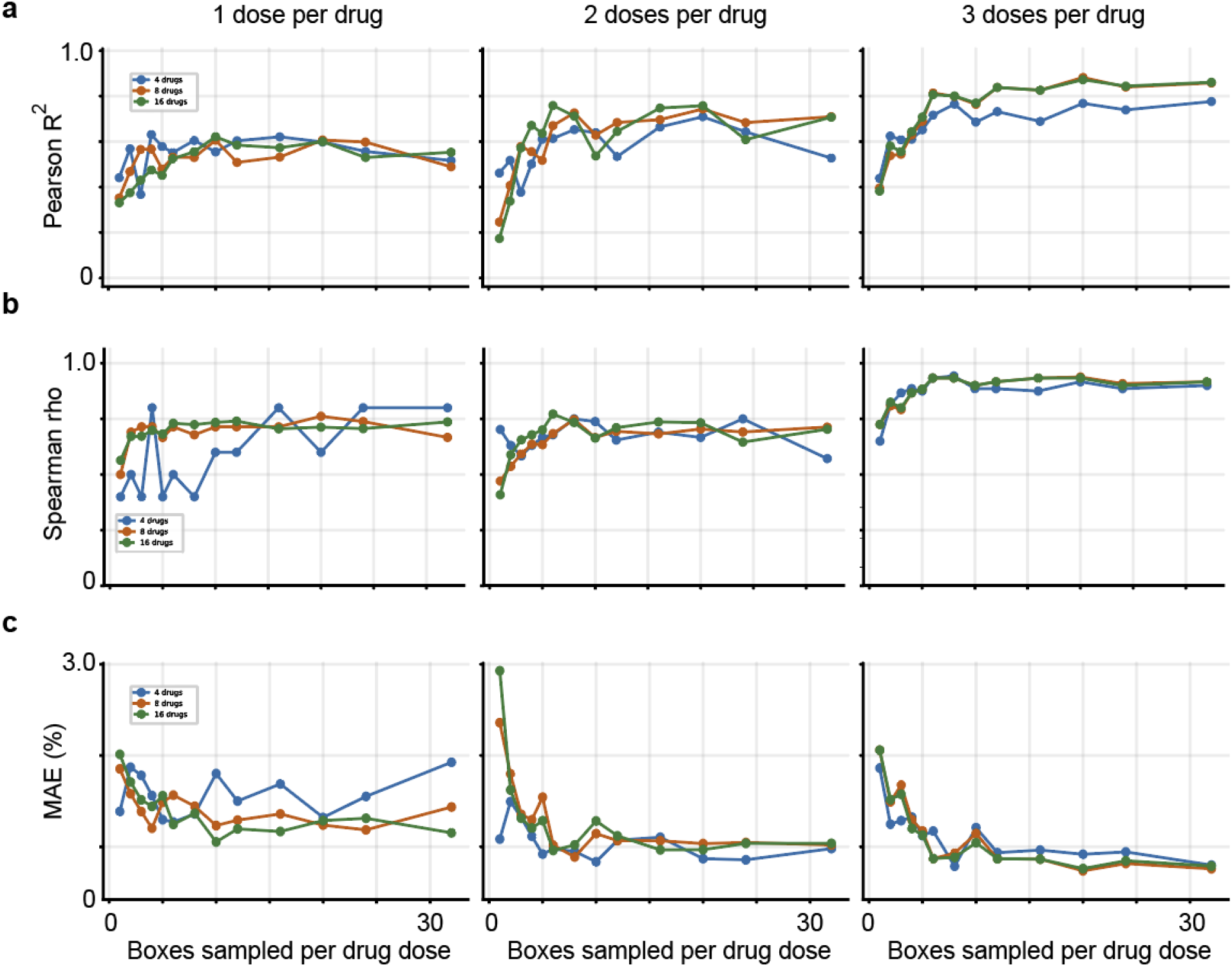
Power analysis. Number of drugs and doses needed for an acceptable Pearson R^2 (a), Spearman rho (b), and median absolute error (MAE) (c).

